# TRAP1 and cyclophilin D compete at OSCP subunit to regulate enzymatic activity and permeability transition pore opening by F-ATP synthase

**DOI:** 10.1101/2021.12.06.471412

**Authors:** Giuseppe Cannino, Andrea Urbani, Marco Gaspari, Mariaconcetta Varano, Alessandro Negro, Antonio Filippi, Francesco Ciscato, Ionica Masgras, Christoph Gerle, Elena Tibaldi, Anna Maria Brunati, Giovanna Lippe, Paolo Bernardi, Andrea Rasola

## Abstract

Binding of the mitochondrial chaperone TRAP1 to client proteins shapes cell bioenergetic and proteostatic adaptations, but the panel of TRAP1 clients is only partially defined. Here we show that TRAP1 interacts with F-ATP synthase, the protein complex that provides most cellular ATP. TRAP1 competes with the peptidyl-prolyl *cis*-*trans* isomerase cyclophilin D (CyPD) for binding to the oligomycin sensitivity-conferring protein (OSCP) subunit of F-ATP synthase, increasing its catalytic activity and counteracting the inhibitory effect of CyPD. Moreover, TRAP1 inhibits opening of the permeability transition pore (PTP) formed by F-ATP synthase and effectively antagonizes the PTP-inducing effect of CyPD, which elicits mitochondrial depolarization and cell death. Consistently, electrophysiological measurements indicate that TRAP1 and CyPD compete in the modulation of channel activity of purified F-ATP synthase, resulting in PTP inhibition and activation, respectively, and outcompeting each other effect on the channel. Moreover, TRAP1 counteracts PTP induction by CyPD, whereas CyPD reverses TRAP1-mediated PTP inhibition. Our data identify TRAP1 as a F-ATP synthase regulator that can influence cell bioenergetics and survival and can be targeted in pathological conditions where these processes are dysregulated, such as cancer.

## Introduction

F-ATP synthase is the rotary nanomachine that synthesizes most cellular ATP by utilizing the H^+^ gradient created by H^+^ pumping coupled to electron transport across the inner membrane [1–3]. Several pieces of evidence indicate that F-ATP synthase can also form a mitochondrial channel, the permeability transition pore (PTP) [4–6]. Indeed, the electrophysiological and pharmacological features of a channel formed by highly purified F-ATP synthase [6] are superimposable to those of the mitochondrial megachannel previously identified in mitoplasts as the PTP [7], and ablation of specific F-ATP synthase subunits affects its conductance [8]. Prolonged PTP openings prompt mitochondrial depolarization and commit cells to death following a variety of stress stimuli [9, 10]. Therefore, the F-ATP synthase is at the crossroad of cell survival and death, suggesting that it might be controlled by complex regulatory circuitries.

The mitochondrial peptidyl-prolyl *cis-trans* isomerase cyclophilin D (CyPD) is a well-known proteinaceous regulator of the F-ATP synthase. CyPD binding to OSCP, a component of the peripheral stalk of the holoenzyme, results in the partial inhibition of F-ATP synthase enzymatic activity [11] and in the sensitization to PTP opening [12]. CyPD also interacts with the mitochondrial paralog of the HSP90 chaperone family TRAP1, an important bioenergetic regulator [13] that was proposed to exert an inhibitory role on PTP opening [14–16]. TRAP1 down-modulates oxidative phosphorylation (OXPHOS) by inhibiting cytochrome *c* oxidase and succinate dehydrogenase (SDH) [17, 18] and can establish a pseudohypoxic phenotype mediated by succinate-dependent induction of the HIF1 transcriptional program [17]. In turn, HIF1 activation increases TRAP1 expression, and TRAP1 is crucial in maintaining a low oxygen consumption rate under hypoxia [19]. These metabolic changes could tune metabolic adaptation to changing environmental conditions such as those experienced during tumor growth. Accordingly, genetic or pharmacological TRAP1 inhibition hampers tumorigenicity in several models of neoplastic progression [20–22], whereas TRAP1 induction correlates with drug resistance, progression and metastatic spreading in a number of malignancies [23]. Nonetheless, TRAP1 expression is downregulated in some specific tumor types and stages [24] suggesting that its contribution to tumor growth could be context-dependent.

These multifaceted effects of TRAP1 could be explained by the engagement of different client subsets finely pitched by post-translational modifications (PTMs) [18,25–28] in multimeric protein assemblies [29, 30], resulting in the modulation of diverse biochemical circuits. A recent survey identified several putative TRAP1 clients, including subunits of F-ATP synthase [29], even if a thorough characterization of such interactions and of their functional effects is still lacking. Here, we find that TRAP1 competes with CyPD for binding to OSCP and increases F-ATP synthase enzymatic activity while decreasing its channel activity. Our data support a model in which TRAP1 and CyPD influence bioenergetic features and survival to noxious conditions of cells by mutually binding to F-ATP synthase, with important implications in pathophysiological conditions such as neoplastic transformation or adaptations to hypoxia.

## Results and Discussion

### TRAP1 interacts with F**-**ATP synthase and CyPD

We performed an unbiased nanoLC-MS/MS mass spectrometry analysis to search for TRAP1 interactors in co-immunoprecipitation experiments on human U87 glioblastoma cells, where TRAP1 displays a pro-neoplastic activity [25]. We found 95 potential TRAP1 mitochondrial interactors, including seven F-ATP synthase subunits (F_1_ sector subunits α, β and γ, peripheral stalk subunits OSCP, d and b, and membrane subunit g) [5, 31] as well as the F-ATP synthase regulator CyPD (Figs 1A, 1B, EV1 and EV2 and Tables EV1 and EV2). Overlay assays confirmed the binding of TRAP1 both to the α, β, γ, b, OSCP and g subunits (Fig 1C) of a highly purified F-ATP synthase preparation from bovine heart [32], and to CyPD (Fig 1D). In a mirror experiment, purified TRAP1 [20] was found to bind CyPD (Fig 1E), in accord with previous data [15, 16]. A direct interaction between TRAP1 and CyPD was also explored with surface plasmon resonance (SPR) (Fig 1F). CyPD interacts with F-ATP synthase subunit OSCP [11,33–35], which is placed on top of the “crown” region of the holoenzyme and connects the peripheral stalk with the catalytic portion of F_1_ [5]. SPR experiments confirmed CyPD binding to OSCP (Fig EV2A) and revealed a direct interaction between TRAP1 and OSCP (Fig 1G). Gel filtration chromatography analyses on U87 mitochondria showed co-localization between TRAP1 and F-ATP synthase subunits in high molecular weight fractions (Fig 1H). Moreover, TRAP1 and CyPD associated with F-ATP synthase both in its monomeric and dimeric form (Fig 1I), the latter constituting the physiological unit of the holoenzyme [31].

**Figure 1.**
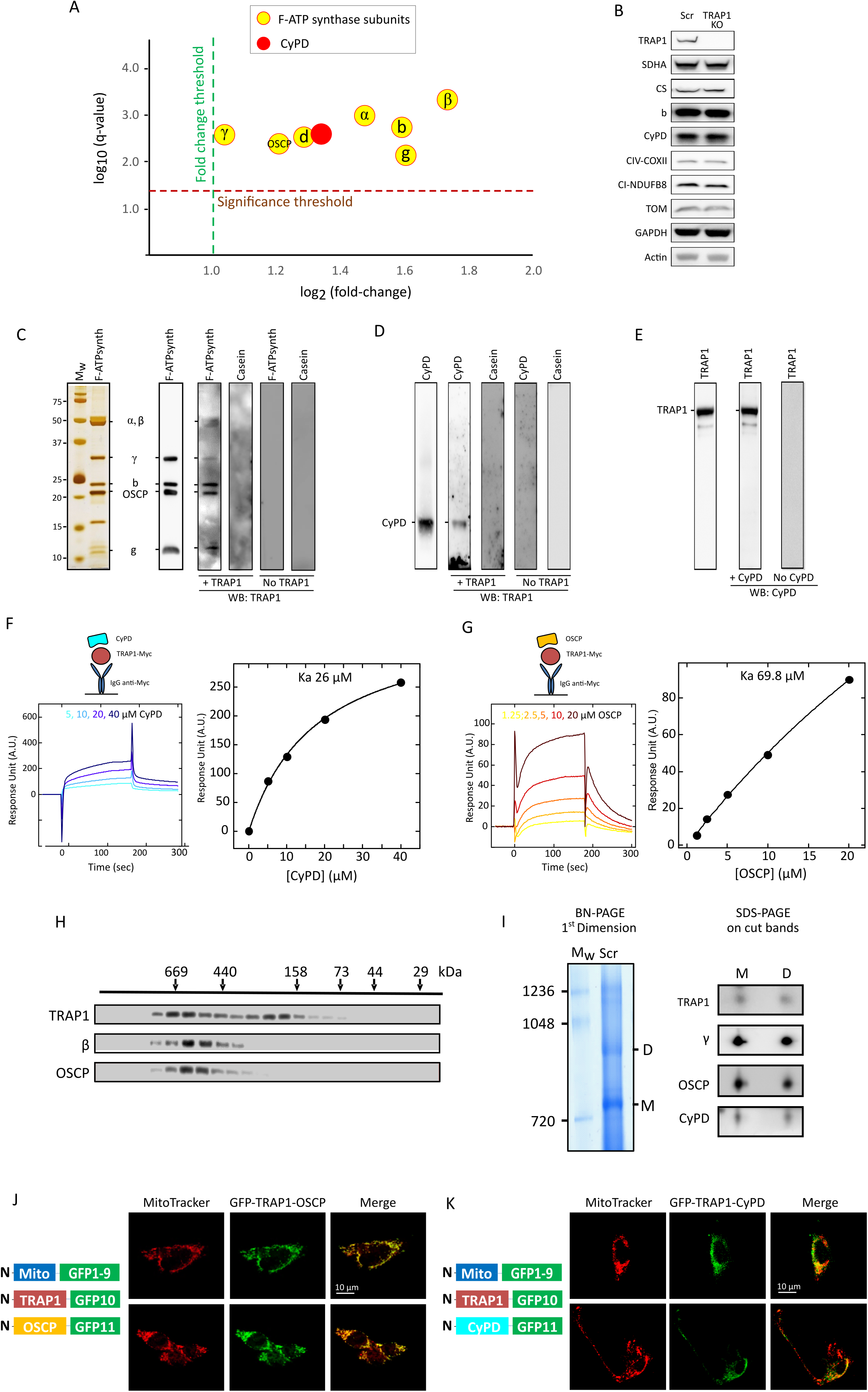
TRAP1 interacts with F-ATP synthase subunit OSCP and with CyPD. A NanoLC-MS/MS mass spectrometry analysis of TRAP1 interactors performed on mitochondria from U87 cells identifies F-ATP synthase subunits and CyPD. B Western immunoblot analysis of mitochondrial markers in human U87 glioblastoma cells with or without TRAP1 (infected with a vector containing a scrambled sequence, SCR, or with a knock-out one, KO, respectively). C-E Overlay assays. In (C), silver staining (left) and Western immunoblot for purified F-ATP synthase (center) were carried out on enzyme subunits separated by SDS-PAGE. Samples transferred onto nitrocellulose were then incubated with or without the purified TRAP1 protein and binding was detected with an anti-TRAP1 antibody. In (D) Western immunoblot on purified CyPD followed by sample transfer onto nitrocellulose, incubation with or without the purified TRAP1 protein and detection with anti-TRAP1 antibody; in (E) Western immunoblot on purified TRAP1 followed by sample transfer onto nitrocellulose, incubation with or without the purified CyPD protein and detection with anti-CyPD antibody. Casein was used as a negative control in all overlay assays. F, G SPR analysis of the binding between CyPD and TRAP1 (F) and between OSCP and TRAP1 (G). Representative schemes of the SPR procedure followed to measure CyPD or OSCP binding to Myc-TRAP1 are shown in the upper left part of each figure. Representative sensorgrams obtained after the injection of the reported concentrations of CyPD or OSCP over Myc-TRAP1, previously immobilized on the surface of the CM5 sensor chips, are reported in the lower left part of each figure. The average response from 0-180 s was plotted as a function of CyPD/OSCP concentration and fitted to a Langmuir interaction model to determine K_a_ (right part of each figure). H Gel filtration chromatography analysis of TRAP1 distribution in mitochondria of human U87 glioblastoma cells. OSCP and β subunits were used to assess the presence of F-ATP synthase. I BN-PAGE performed on isolated mitochondria (left) showing F-ATP synthase monomers and dimers (M and D, respectively). M and D bands were cut, run on SDS-PAGE and probed for the presence of F-ATP synthase subunits γ and OSCP, TRAP1 and CyPD. J, K Tripartite split GFP experiment on sMPNST cells co-transfected with GFP1–9 fused to a mitochondrial import sequence, together with GFP10-TRAP1 and GFP11-OSCP (J), or with GFP10-TRAP1 and GFP11-CyPD (K). Green dots indicate mitochondrial interaction between TRAP1 and OSCP (J) or TRAP1 and CyPD (K). MitoTracker (red dots) was used to stain mitochondria; superimposition of MitoTracker and GFP signal is shown as yellow dots and indicates that protein interaction occurs in mitochondria.

We then addressed whether these protein-protein interactions occur *in situ* by using a split-GFP approach [21, 36]. We co-expressed (1) a GFP portion (GFP1–9) fused with a mitochondrial import sequence; (2) a GFP fragment (GFP10) fused to one putative interactor; and (3) the final GFP moiety, GFP11, associated to the second potential interactor. GFP reconstitution indicates a direct interaction of the three GFP portions and therefore of the two proteins of interest in mitochondria. We carried out these tripartite split-GFP experiments on malignant peripheral nerve sheath tumor (sMPNST) cells, where TRAP1 is pro-tumorigenic [20, 21]. The green signal observed in cells co-expressing Mito-GFP1-9, CyPD-GFP10 and OSCP-GFP11, and its merging with the red staining by the mitochondrial probe MitoTracker Red, confirm the expected binding between CyPD and OSCP (Fig EV2B). The same approach was carried out on cells co-expressing Mito-GFP1-9 with either TRAP1-GFP10 and OSCP-GFP11 or TRAP1-GFP10 and CyPD-GFP11, revealing a direct mitochondrial interaction between OSCP and TRAP1 (Fig 1J) and between CyPD and TRAP1 (Fig 1K), respectively.

Altogether, these data indicate that TRAP1 interacts with F-ATP synthase and CyPD in mitochondria and suggest the presence of multimeric platforms that could confer regulatory flexibility to F-ATP synthase activity.

### TRAP1 affects the binding of CyPD to OSCP

The CyPD/OSCP interaction mildly inhibits ATP synthesis and hydrolysis, and sensitizes the PTP to opening [37, 38]. We therefore evaluated whether TRAP1 regulates F-ATP synthase enzymatic activity and PTP opening through its interactions with CyPD and OSCP. Gel filtration chromatography experiments detected CyPD, OSCP and TRAP1 in the same high molecular weight fractions (Fig 2A). In TRAP1 knock-out cells all CyPD co-purified with OSCP in these fractions, whereas in TRAP1-expressing cells a low molecular weight CyPD signal was also detected, corresponding to a portion of CyPD unbound to protein complexes (Fig 2A). Moreover, the amount of CyPD that co-immunoprecipitated with OSCP (Fig 2B) or co-migrated with F-ATP synthase dimers and monomers (Fig 2C) was higher in the absence of TRAP1, suggesting competition for binding. In keeping with the ability of TRAP1 to disrupt the interaction of CyPD with OSCP, split GFP experiments showed that the binding between CyPD and OSCP was enhanced by knocking-out TRAP1 or by targeting it with the highly selective TRAP1 inhibitor i5 [21] (Figs 2D and 2E). Treatment with i5 also decreased the binding of TRAP1 to both OSCP and CyPD (Figs EV3A and EV3B). Consistently, both knocking-out CyPD and treatment with cyclosporin A (CsA), which displaces CyPD from F-ATP synthase [11], increased the interaction of TRAP1 with OSCP (Figs 2F and 2G) and inhibited CyPD/OSCP binding (Fig EV3C). These data indicate that TRAP1 and CyPD compete for the binding to OSCP and suggest that TRAP1 could counteract both the inhibitory effect of CyPD on the enzymatic activity of F-ATP synthase and its promoting effect on PTP opening.

**Figure 2.**
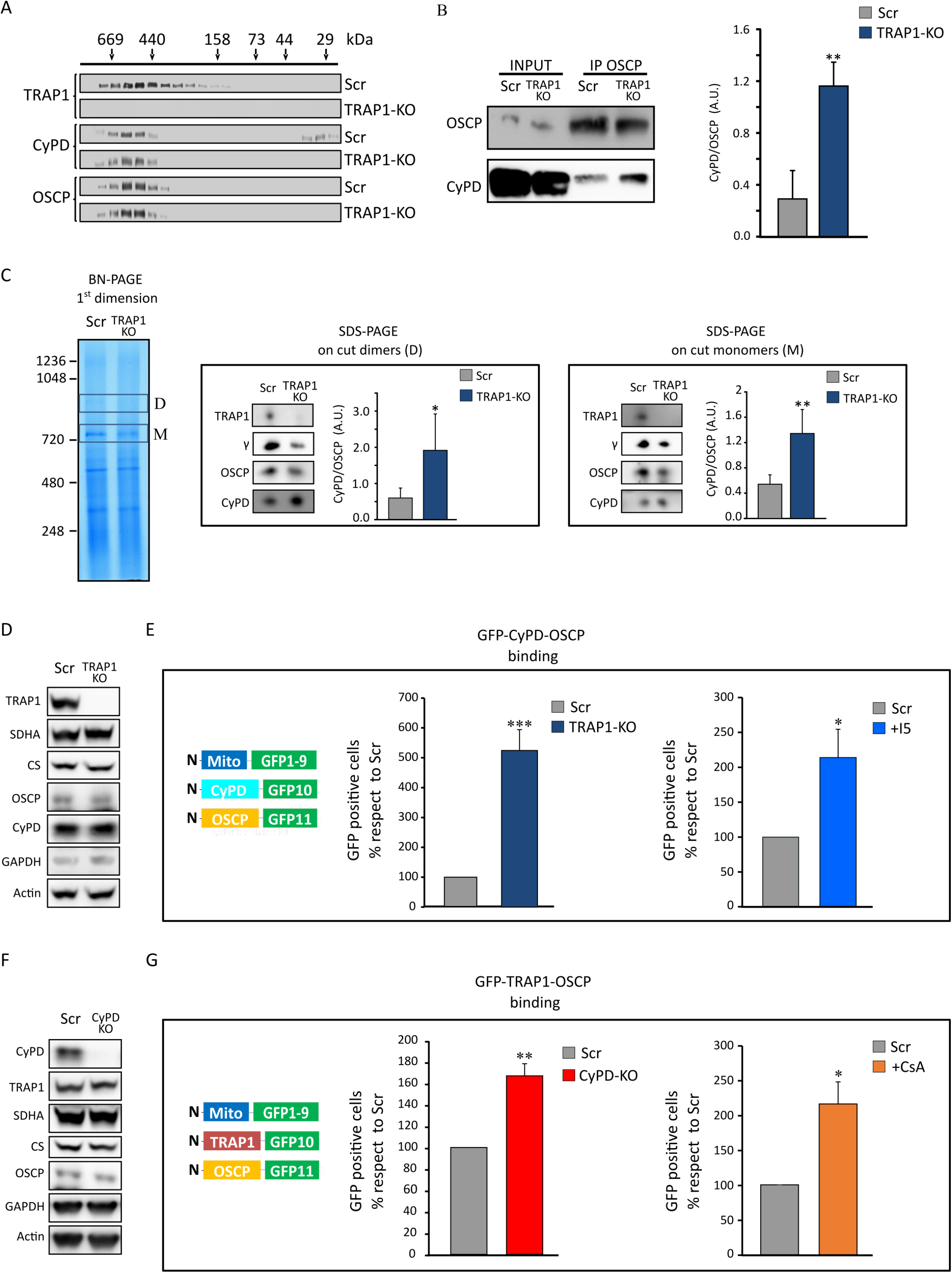
TRAP1 and CyPD compete for interaction with OSCP. A Gel filtration chromatography analysis of CyPD and OSCP distribution in mitochondria of human U87 glioblastoma cells with or without TRAP1 (SCR and KO, respectively). B Immunoprecipitation of OSCP from mitochondria of U87 cells. On the right, quantification of the amount of CyPD bound to OSCP. Data are presented as mean ± SEM of 4 independent experiments; **p < 0.01 with a paired two-tailed Student’s *t* test. C BN-PAGE and Western immunoblotting of F-ATP synthase extracted from mitochondria of U87 cells. D (dimer) and M (monomer) bands were cut, run on SDS-PAGE and probed with antibodies for TRAP1, CyPD and F-ATP synthase subunits γ and OSCP. Histograms refer to the mean ± SEM of CyPD bound to dimers (center) or monomers (right) normalized for the content of OSCP subunit of 4 independent experiments. D, F Western immunoblots of mitochondrial markers from sMPNST cells with or without TRAP1 (D; SCR and KO, respectively), and with or without CyPD (F; SCR and KO, respectively); actin was used as a loading control. E, G Tripartite split GFP experiment on sMPNST cells co-transfected with GFP1–9 fused to a mitochondrial import sequence, together with GFP10-CyPD and GFP11-OSCP (E), or with GFP10-TRAP1 and GFP11-OSCP (G). Experiment analysis was carried out as in Figure 1J, K. Quantification of GFP-positive cells was carried out without or with treatment with 100µM TRAP1 inhibitor i5 (histograms in E; cells analyzed >1300) or without or with treatment with 2µM CsA (histograms in G; cells analyzed >1500). Data are presented as mean ± SEM. ^∗∗^p< 0.01 with a paired two-tailed Student’s *t* test.

### TRAP1 increases F-ATP synthase enzymatic activity

We then explored the functional effects of TRAP1 binding to F-ATP synthase. Both ablation of TRAP1 and treatment with i5 inhibited the hydrolytic activity of F-ATP synthase in U87 cells (Fig 3A), and knocking-out TRAP1 markedly decreased the in-gel hydrolytic activity of F-ATP synthase dimers without affecting the dimer/monomer protein levels (Fig 3B). Incubation with TRAP1 also increased the hydrolytic activity of F-ATP synthase in submitochondrial particles where endogenous CyPD had been removed, and this induction was blunted by treatment with i5. In keeping with previous reports, CyPD had the opposite effect (*i.e.* it decreased the enzymatic activity in a CsA-sensitive way) [11], and we found that TRAP1 abrogated the inhibitory effect of CyPD (Fig 3C). These observations are consistent with a model in which TRAP1 competes with CyPD for OSCP binding, enhancing the ATPase activity of F-ATP synthase.

**Figure 3.**
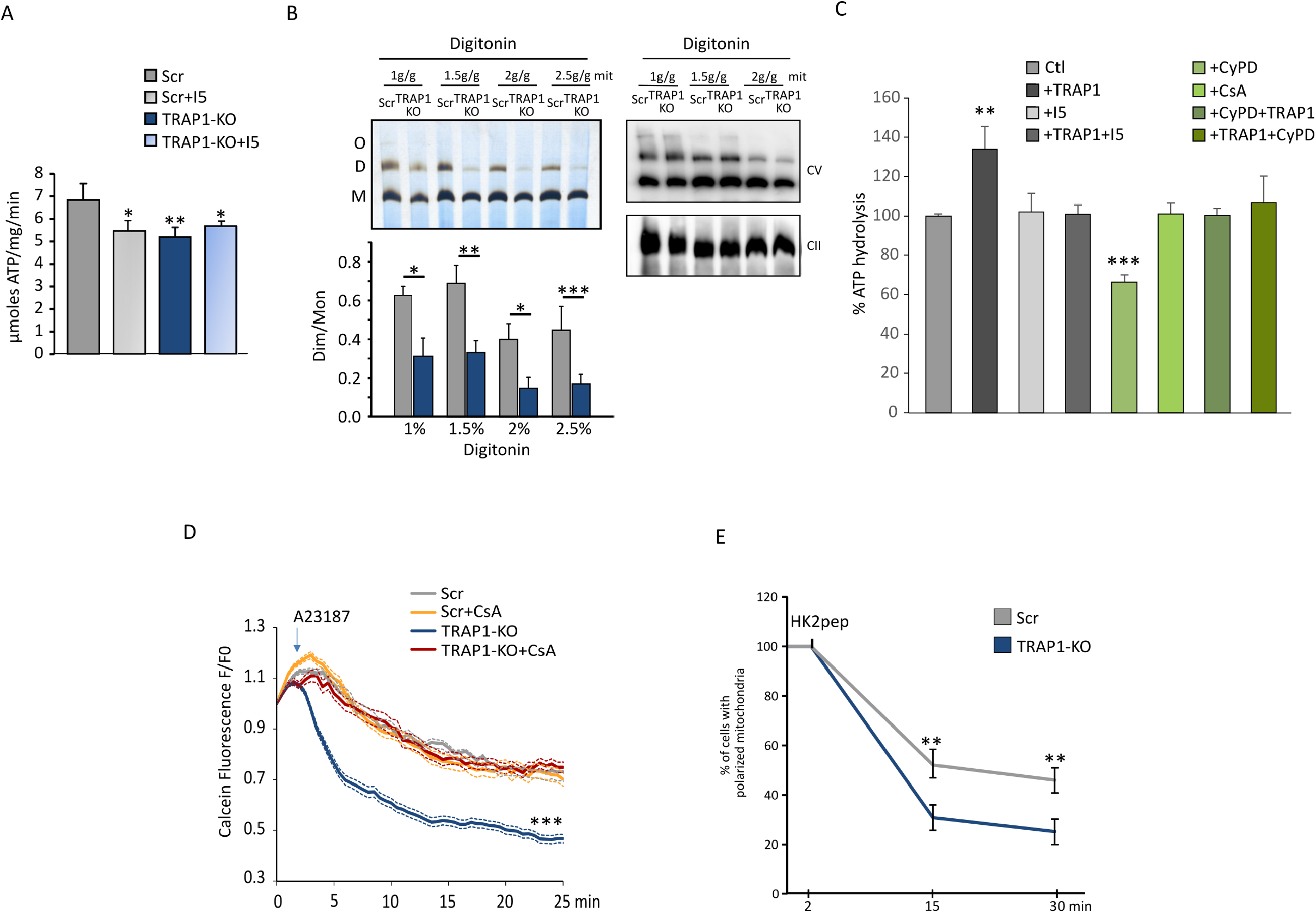
Effect of TRAP1 interaction on F-ATP synthase enzymatic activity and PTP opening. A Oligomycin-sensitive ATPase activity on human U87 glioblastoma cells with or without TRAP1 (SCR and KO, respectively). Where indicated, the TRAP1 inhibitor i5 was used at 25 µM. The histogram reports the mean ± SEM of 4 independent experiments. B BN-PAGE followed by in gel ATPase activity (top left) on U87 mitochondria extracted at the indicated concentrations of digitonin. Quantification of ATPase activity is reported as the mean dimer/monomer ratio from three independent measurements; *p 0.01, **p 0.004 (bottom, left). BN-PAGE followed by Western immunoblotting (top, right) was probed with antibodies for F-ATP synthase subunit β to detect enzyme monomers, dimers and oligomers; anti-SDHA antibody was used as a loading control. C ATPase activity measured in submitochondrial particles (SMPs) from pig heart pre- incubated with the indicated amounts of TRAP1 or CyPD alone or sequentially for 5 min at 37°C. Where indicated, the TRAP1 inhibitor i5 (100 μM) or CsA (4 μM) were preincubated for 5 min at 37°C. Data are shown as percentage changes with respect to untreated SMPs. D Calcein cobalt assay of PTP opening on U87 cells. Cells were stained with calcein AM at time zero; 2 μM Ca^2+^ ionophore A23187 was added where indicated in the absence or in the presence of 2 μM CsA. Image frames were collected at 30-s intervals and fluorescence values (mean ± SEM n=4) were quantified. E Mitochondrial membrane potential assessment in U87 cells after the addition of 2 µM cl- HK2pep. Cells were stained with TMRM probe (20 nM + 1 µM CsH); data are reported as mean ± SD; **p<0.01 with a TwoWay Anova p<0,001, Bonferroni post-test analysis.

### Ablation of TRAP1 sensitizes PTP opening

By using a calcein/Co^2+^ assay [39], we found that TRAP1 ablation sensitizes U87 cells to PTP opening following treatment with the Ca^2+^ ionophore A23187 (Fig 3D). CyPD inhibition by CsA desensitized pore opening in TRAP1 knock-out cells, while it was ineffective in their wild-type counterparts (Fig 3D), in accord with a competition between TRAP1 and CyPD for binding to F- ATP synthase. Prolonged PTP openings cause mitochondrial membrane depolarization and eventually cell death [9]. In keeping with an inhibitory role of TRAP1 on PTP opening, we observed that its ablation accelerates mitochondrial depolarization after cell treatment with cl- HK2pep (Fig 3E), a peptide that induces Ca^2+^-dependent PTP opening, mitochondrial depolarization and cell death by displacing hexokinase 2 from endoplasmic reticulum/mitochondria contact sites [40].

We directly assessed the effect of TRAP1 on PTP opening by using a highly purified F-ATP synthase, which is able to elicit channel activity matching the electrophysiological features of the PTP after insertion in planar lipid bilayers [6]. We observed high-conductance channels with multiple substates upon addition of F-ATP synthase and Ca^2+^ to the lipid bilayers. Channel behavior was mostly flickering and in lower conductance sublevels, as previously observed in the absence of additional PTP inducers, but stable channel openings reaching up to one or more PTP full conductances were also detected (Figs 4A, 4G, 4I, EV4A and EV4B). Addition of TRAP1 in 1:1 to 3:1 molar ratio with F-ATP synthase reduced channel activity in a concentration-dependent way in 8 out of 14 experiments, leading to almost total suppression of the currents in 3 experiments. TRAP1 mostly abolished the higher conductance levels, leaving very low residual currents and sporadic channel openings (Figs 4B, 4C, 4H and EV4C), whereas it had negligible effects on the low-conductance flickering behavior of the channel (Figs 4J and 4K). Residual currents were still sensitive to PTP inhibitors, such as Ba^2+^ (Fig 4D). Further addition of Ca^2+^ could only partially restore channel activity, which was still sensitive to the inhibitory effect of TRAP1 (Figs 4E and 4F).

**Figure 4.**
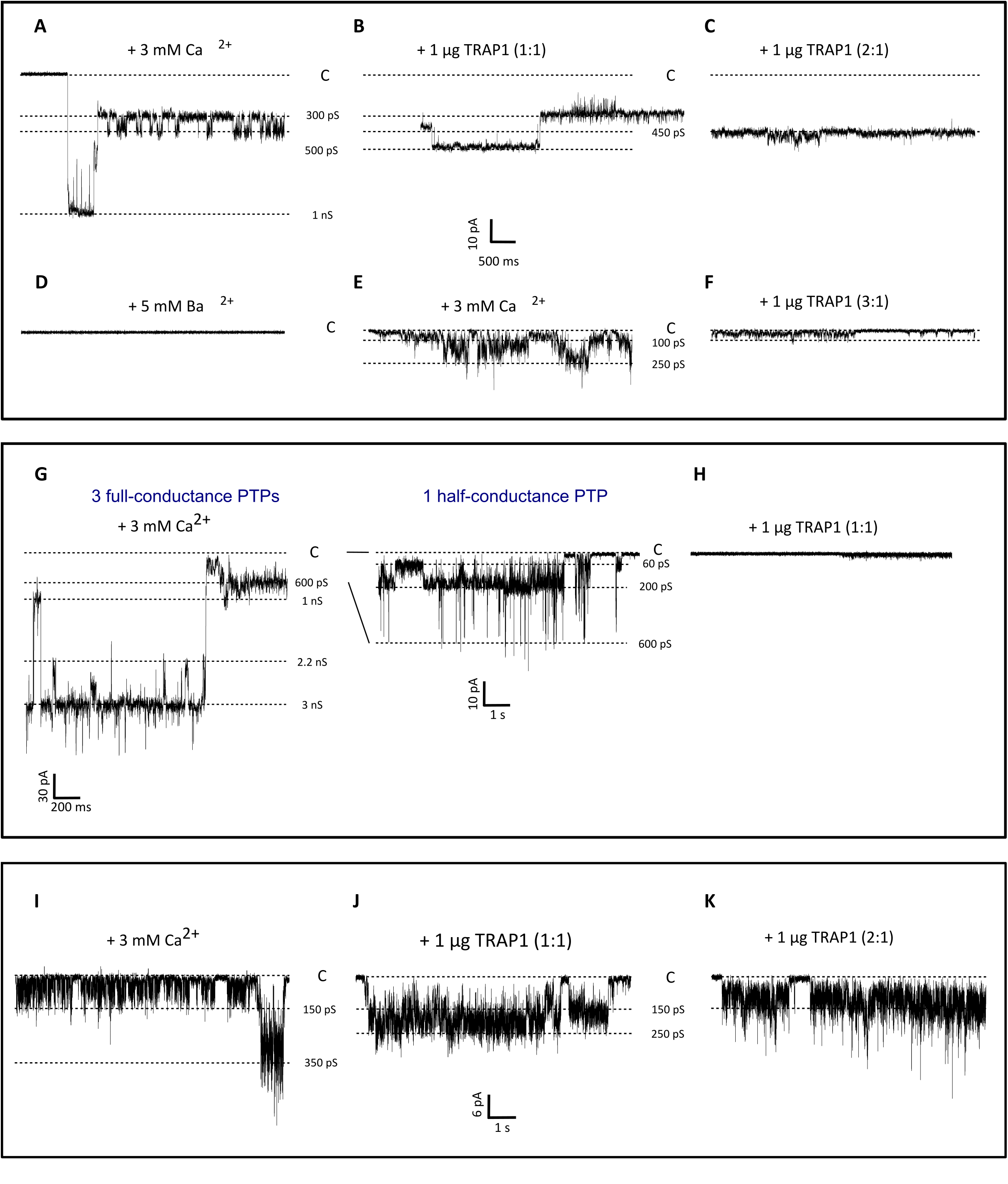
Effect of TRAP1 on F-ATP synthase channel activity. A-F High conductance channel activity, corresponding to one full-conductance PTP, detected after addition to the *cis* side of the bilayer of 8 µg F-ATP synthase in the planar lipid bilayer after addition of 3 mM Ca^2+^ (A; V_cis_ = –60 mV). The effects on channel activity of subsequent additions of TRAP1 (B: molar ratio with F-ATP synthase of 1:1; G_max_ = 500 pS; C: molar ratio with F-ATP synthase of 2:1; G_max_ = 450 pS), 5 mM Ba^2+^ (D), further 3 mM Ca^2+^ (E; G_max_ = 250 pS); further 1 µg TRAP1 (F; molar ratio with F-ATP synthase of 3:1; G_max_ = 100 pS) are reported. G, H Very high and complex channel activity, corresponding to up to three PTPs at full-conductance (unitary conductance 1 nS), detected after addition of 3 mM Ca^2+^ (g; V_cis_ = –60 mV); see Figure EV4 for the whole trace. Current levels corresponding to a single PTP in full-conductance and half-conductance are also visible (1 nS and 600 pS), as well as levels corresponding to other multiple PTP openings (2.2 nS, 3 nS). On the right, channel activity at lower conductance levels is shown at a higher magnification. In (H), addition of TRAP1 (at a molar ratio with F-ATP synthase of 1:1) led to almost total current suppression. Sporadic channel openings are reported in Figure EV4. I-K Flickering channel activity at low conductance detected after addition of 3 mM Ca^2+^. In (I), the most represented conductance level was at 150 pS, while the maximal conductance state was at 350 pS. The effects on channel activity of subsequent additions of TRAP1 (J: molar ratio with F-ATP synthase of 1:1; G_max_ = 250 pS; K: molar ratio with F-ATP synthase of 2:1; G_max_ = 250 pS) are reported. All along the figure, the prevalent conductance levels are marked with dotted lines and the conductance values are reported; the closed state is denoted with C.

We also studied the effect of CyPD on the currents elicited by F-ATP synthase, and assessed a possible functional interplay between CyPD and TRAP1. We found that CyPD, in a molar ratio of 2:1 with F-ATP synthase, increased Ca^2+^-induced currents to high-conductance levels compatible with one or more PTPs (Figs 5A and 5B). The subsequent addition of TRAP1 in a molar ratio of 1:1 with F-ATP synthase reduced channel activity (Fig 5C), even if high conductance openings could still be observed (Fig EV5). A further addition of TRAP1 to reach a molar ratio of 2:1 with F-ATP synthase stabilized the lower conductance states, reducing the amplitude of the remaining high-conductance channel openings (Figs 5D and EV5B). In a mirror experiment, addition of CyPD could restore Ca^2+^-induced, high-conductance F-ATP synthase currents that had been reduced by TRAP1 (molar ratio of 2:1 with F-ATP synthase for both CyPD and TRAP1) (Figs 5E-G). A further addition of CyPD up to a 5:1 molar ratio with F-ATP led to flickering openings to conductance levels compatible with one or more fully open PTPs (Fig 5H). These data indicate that TRAP1 and CyPD compete to modulate opening of the PTP channel with opposite functional effects.

**Figure 5.**
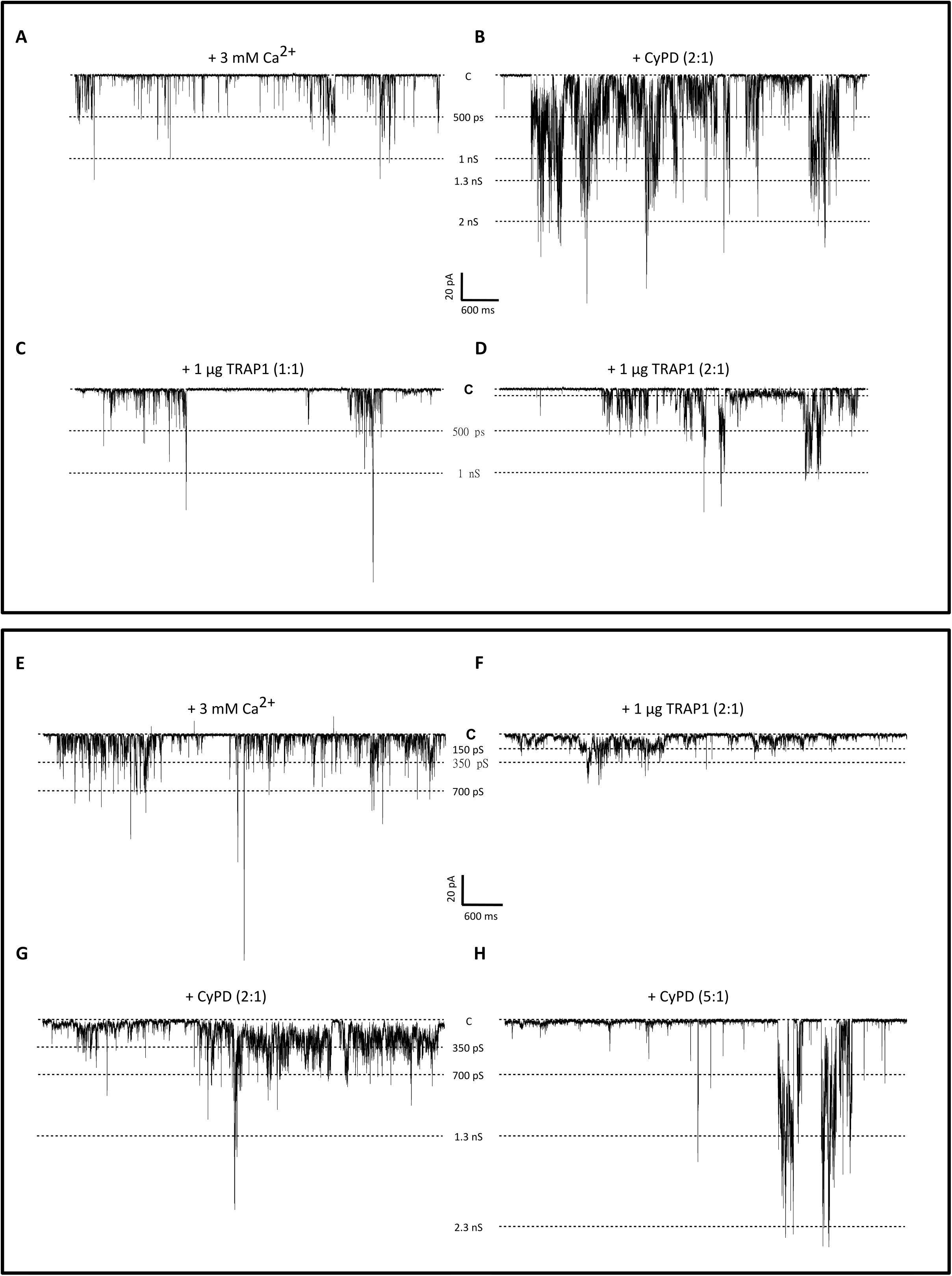
Opposite effects and competition of TRAP1 and CyPD on F-ATP synthase channel activity. A-D High conductance, flickering channel activity detected after direct incorporation of 16 µg F- ATP synthase in the planar lipid bilayer and addition of 3 mM Ca^2+^ (A; V_cis_ = –60 mV). The effects on channel activity of subsequent additions of CyPD (B; molar ratio with F-ATP synthase of 2:1) and TRAP1 (C: molar ratio with F-ATP synthase of 1:1; D: molar ratio with F-ATP synthase of 2:1) are reported. E-H High conductance, flickering, channel activity detected in the presence of 3 mM Ca^2+^ (E; V_cis_ = –60 mV). The effects on channel activity of subsequent additions of TRAP1 (F; molar ratio with F-ATP synthase of 2:1) and CyPD (G: molar ratio with F-ATP synthase of 2:1; H: molar ratio with F-ATP synthase of 5:1) are reported. In all figure panels, the prevalent conductance levels are marked with dotted lines and the conductance values are indicated; the closed state is denoted with C. Either TRAP1 or CyPD were added at increasing concentrations after Ca^2+^-induced channel activity was detected.

In the current study, we document the interaction between the molecular chaperone TRAP1, the F- ATP synthase and the peptidyl-prolyl *cis-trans* isomerase CyPD. We demonstrate that TRAP1 enhances the enzymatic activity of F-ATP synthase and abrogates the inhibitory effect of CyPD. Moreover, we find that TRAP1 antagonizes PTP channel openings observed with purified F-ATP synthase, whereas CyPD enhances them, and that TRAP1 and CyPD outcompete each other for PTP regulation.

The F-ATP synthase holoenzyme synthesizes most cellular ATP via a mechanism of rotary catalysis. Its subunits are arranged in an F_O_ sector embedded in the inner mitochondrial membrane, through which protons generated by respiratory complexes translocate to promote rotation, conveyed into the matrix where ATP is produced or hydrolyzed by the catalytic F_1_ sector. OSCP, together with F_6_, d, A6L and b subunits, forms the enzyme peripheral stalk. This acts as a stator that prevents co-rotation of F_1_ with F_O_ and connects the two sectors through subunits e, f, g, a, j and k, which also form the dimerization interface. Central stalk components of F_1_ (γ, δ, and ε subunits) connect the membrane ring formed by c subunits to a α_3_β_3_ hexamer in F_1_ head, and ATP synthesis occurs at the interfaces between catalytic β subunits and non-catalytic α subunits. [5,41,42]. Among potential TRAP1 interactors, we and others [29] have found the F_1_ components α, β and γ and the peripheral stalk subunit b, g, d and OSCP, suggesting the possibility of a multifaceted regulation of the enzyme functions. The interaction between TRAP1 and OSCP might be particularly important for F-ATP synthase regulation. OSCP couples the peripheral stalk to the “crown” at the head of the F_1_ sector, clamping it in the correct position [41] but allowing a number of conformations during catalysis; thus, OSCP is placed in a strategical position for interacting with regulatory components of F-ATP synthase [42]. Accordingly, OSCP interacts with: (i) CyPD in a CsA-sensitive way, decreasing enzyme activity and sensitizing PTP to opening [11]; (ii) the benzodiazepine Bz-423, which mimics the effects of CyPD on both enzyme activity and PTP opening [4]; (iii) the tumor suppressor p53 during F-ATP synthase assembly [43], with p53 inducing PTP opening following CyPD binding [44] and displacing CyPD/TRAP1 interaction *in vitro* [16]; and (iv) the NAD^+^-dependent deacetylase SIRT3 [35], which decreases CyPD binding by de-acetylation of OSCP Lys70 [34]; and (v) estrogen receptor β in competition with CyPD, which results in cytoprotection [33]. Moreover, OSCP Cys141 allows CyPD shielding from PTP induction by oxidants [45], whereas OSCP His112 mediates PTP inhibition by protons, and His112 mutants form channels insensitive to inhibition by acidic pH [46]. A recent cryo-EM mapping of F-ATP synthase suggests a “death finger” model [47] for PTP induction, whereby CyPD binding to OSCP in the presence of Ca^2+^, would transmit a conformational change along the peripheral stalk to the e subunit, which would retract from the c ring, thus pulling out the outer lipid plug causing channel opening [5].

Our observations that TRAP1 binds to OSCP in competition with CyPD, counteracting the inducing effects of the latter on F-ATP synthase, add an important element to this complex picture. TRAP1 could either outcompete the CyPD/OSCP interaction or sequester CyPD away from F-ATP synthase, two non-mutually-exclusive options. We propose that OSCP acts as a hub that finely tunes both the enzymatic activity and the conversion of F-ATP synthase to the PTP by interacting with different protein regulators. The choice of the OSCP binding partner(s) would result in specific biochemical outcomes, with the effect of optimizing biological outputs to changing environmental conditions. The composition of this molecular platform could be dictated by signal transduction cascades connecting F-ATP synthase with the metabolic needs of the cell or with stress signals. Both TRAP1 and CyPD can undergo many PTMs that affect their activity, with a particular relevance in tumor models. Phosphorylation downstream to oncogenic dysregulation of Ras/ERK signalling inhibits the PTP-sensitizing effects of CyPD [48] and increases the chaperone activity and the pro-neoplastic effects of TRAP1 [25]. Additional PTMs include oxidation, S-nitrosylation, S-palmitoylation, S-glutathionylation and acetylation for CyPD [12], and S-nitrosylation and acetylation for TRAP1, as well as its formation of tetramers [24]. It is easy to see how the combination of these modifications can finely tune access to the F-ATP synthase and ATP synthesis or channel formation activities. It is conceivable that under conditions of nutrient and oxygen shortage, which can occur during embryonic development or in the growing tumor mass, partial suppression of F-ATP synthase activity could contribute to the establishment of an advantageous aerobic glycolysis phenotype, whereby OXPHOS is minimized in favor of glycolysis and/or glutaminolysis. Notably, we observe that neither TRAP1 nor CyPD change the levels of F-ATP synthase monomers or dimers. Hence, chaperone regulation of the enzyme must be achieved through direct protein interaction(s), allowing rapid and flexible control.

Prolonged PTP opening constitutes a point of no return in cell death induction, making its regulation by protein-protein interactions an issue of the utmost interest in the context of a variety of pathophysiological conditions [49]. Here we demonstrate that at least two proteins, CyPD and TRAP1, regulate PTP opening in electrophysiology experiments where a purified F-ATP synthase was reconstituted into lipid bilayers. It should be noted that in the case of CyPD, even if it constitutes the best-characterized proteinaceous PTP regulator, a direct effect on the channel activity of F-ATP synthase was never shown. We have observed that CyPD increases F-ATP synthase channel openings and current amplitude in the presence of Ca^2+^, in perfect keeping with its widely documented function of PTP sensitizer. Conversely, we have documented that TRAP1 reduces Ca^2+^-induced F-ATP synthase channel activity. This inhibition can be either partial, lowering the PTP conductance to sub-maximal levels, or complete, fully ablating the currents. Interestingly, TRAP1 has little if any effect on the flickering, low-conductance mode of PTP activity. These observations suggest that a direct interaction between TRAP1 and F-ATP synthase could stabilize sub-conductance levels of PTP, while inhibiting the full-conductance channel. This dual effect could be related to different conformational states of F-ATP synthase that may separately account for transient and for long-lasting PTP openings [50]. Transient openings would provide mitochondria with a fast Ca^2+^ release channel involved in maintaining mitochondrial and cellular Ca^2+^ homeostasis [51–56], while long-lasting openings would lead to persistent mitochondrial depolarization, matrix swelling, release of apoptogenic proteins and eventually cell death [57]. In further keeping with a model where TRAP1 and CyPD compete for the binding to F- ATP synthase, each protein is able to revert the effect on PTP elicited by the other. It must be stressed that inhibition of F-ATP synthase channel by TRAP1 occurs independently of its chaperone activity, which requires ATP that was not provided in our electrophysiological experiments. Interestingly, it was proposed that in the presence of Ca^2+^ TRAP1 may also act as a holdase keeping clients in specific conformations independently of the canonical chaperone activity [58]. Whether a similar mechanism applies to the mode of interaction with F-ATP synthase is an intriguing possibility that deserves further investigation.

Our data predict the existence of multimeric and dynamic protein platforms that finely regulate F- ATP synthase activity between ATP synthesis and channel formation, an issue that bears on cellular bioenergetics and survival. The finding that mitochondrial interactors of TRAP1 also include TCA cycle enzymes and additional OXPHOS proteins, transporters and chaperones points to exciting directions for future investigations. Given that TRAP1 is expressed in many cancers, disruption of its regulatory interactions by highly selective inhibitors [59] may help develop innovative anti- neoplastic treatments.

## Materials and methods

### Cell and sample preparation

Experiments were performed on human U87 glioblastoma cells and mouse malignant peripheral nerve sheath tumor cells (sMPNST cells). TRAP1 and CyPD expression were knocked out using the lenti CRISPRv2 system [60]. Sequences for the guides used to knock out TRAP1 in U87 cells were previously reported [25]. Sequences for the guides used to knock out CyPD in sMPNST cells are: guide 1 5‘CACCGGCGACCCGTACCTGCAGCGA3’ and 5‘AAACTCGCTGCAGGTACGGGTCGCC; guide 2 5‘CACCGGTACACGAGCGGGTTCCCGG3’ and 5‘AAACCCGGGAACCCGCTCGTGTACC3’; and guide 3 5‘CACCGCCCACGTCCAAGTACACGAG3’ and 5‘AAACCTCGTGTACTTGGACGTGGGC3’.

All guides were generated by using the CRISPR design tool (https://chopchop.cbu.uib.no/results/1571645627074.5044/). Oligonucleotide pairs were annealed and cloned into the transfer plasmid lentiCRISPRv2 (Addgene, https://www.addgene.org/52961/) and co- transfected with the three packaging plasmids pMDLg/pRRE, pRSV-Rev and pMD2.G into HEK 293T cells for viral production. Recombinant virus was collected and used to infect cells by standard methods. Infected cells were selected with 1μg/ml puromycin. U87 cells were grown in Minimum essential medium Eagle (MEM) supplemented with 10% fetal bovine serum, glycine (4mM), pyruvate (1 mM) and non-essential amino acids. sMPNST cells were grown in Dulbecco’s modified Eagle’s medium (DMEM) supplemented with 10% fetal bovine serum. Cytoplasmic fractions were obtained by cells kept on ice for 20min in a lysis buffer containing 150 mM NaCl, 20 mM Tris, 5 mM EDTA-Tris, pH 7.4 with the addition of 1% Triton X-100, 10% glycerol and protease /phosphatase inhibitor cocktail (Merck, Darmstadt, Germany) and cleared by centrifugation at 18.000 *g* for 20 min, 4°C. Mitochondria were isolated after cell disruption with a glass-Teflon potter in a buffer composed of 250 mM sucrose, 10 mM Tris-HCl, 0.1 mM EGTA-Tris, pH 7.4.

### Immunoprecipitation

Mitochondria were solubilized on ice for 30 min in lysis buffer and cleared by centrifugation at 18.000 *g* for 20 min, 4 °C. The supernatant was incubated overnight at 4 °C with beads (Invitrogen) previously conjugated with 1 µg of mouse monoclonal anti-human TRAP1 (sc-73604) or mouse monoclonal anti ATP50/OSCP (ab110276) antibodies. Complexes bound to beads were digested according to Bernaudo et al. [61]. Briefly, on-beads digestion was achieved by adding 200 ng trypsin in phosphate buffer (10 min at 37 °C). Alternatively, proteins were eluted with sample buffer containing 2% SDS, 50 mM Tris pH 6.8, 10 % glycerol and 0,00004% Bromophenol blue (5 min at 90 °C) and subjected to Western blotting.

### LC-MS/MS

Partially digested samples were evaporated in speed vacuum and resuspended in 50 µL of HPLC water, in-solution digested and purified by C18 StageTips as described [61]. Lyophilized digests were resuspended in 29 µL of 100 mM TEAB buffer and labelled with tandem mass tags (TMT 10- plex) reagents as described below. A reference sample was created by pooling 4 µL aliquots of each sample. TMT reagents were resuspended in 100 µL of anhydrous acetonitrile; a 10 µL aliquot of each tag solution was employed to label 25 µL of each peptide mixture (one reference pool, four replicates of control samples, four replicates of TRAP1 samples). Labelled peptides were combined, fractionated by SCX StageTips and analyzed by nanoscale liquid chromatography coupled to tandem mass spectrometry as described [61]. Database search was also performed as described in the cited reference, except for the following modification: ^18^O was not indicated as variable modification, whereas TMT 10-plex labelling was set as fixed modification for lysine and N-terminal amino groups. Protein relative quantification was also performed in Proteome Discoverer by relying on TMT mass reporter ions. Protein hits with a minimum of two peptides identified with 95% confidence and an Xcorr of 2 were retained. Protein ratios were calculated relative to the reference sample and normalized on the basis of the content of the bait protein. Thus, the normalization factor for all protein quantification values in a particular sample was Tx/Tm, where Tx was the relative quantification value of TRAP1 for sample x and Tm was the mean relative quantification value of TRAP1 in the four sample replicates. Protein quantification and assessment of significance was determined in Prism 6.0 as described [62].

### Gel filtration chromatography

Cells were disrupted on ice by sonication (2 cycles of 5 sec at 22 Hz with intervals of 15 seconds) in isotonic buffer (50 mM Tris/HCl, pH 7.5, 0.25 M sucrose, 1 mM EDTA and protease and phosphatase inhibitor cocktails). The homogenate was centrifuged for 5 min at 900 *g* and the precipitate (nuclei and unbroken cells) was discarded. The supernatant was centrifuged at 10000 *g* for 20 min. The pellet was resuspended in homogenization buffer and overlaid on cushions of 50 mM Tris/HCl, pH 7.5, containing 1 mM magnesium acetate and 1.25 M sucrose, and centrifuged at 90000 *g* for 60 min using a rotor of the swinging-bucket type. The Golgi apparatus fraction was collected at the 0.25 M/1.25 M sucrose-homogenate interface, while the mitochondria fraction was recovered in the 1.25 M sucrose phase. Mitochondrial fraction was lysed in buffer containing digitonin 0.7%, 20 mM Tris-HCl, pH 7.5, 2 mM EGTA, 150 mM NaCl, protease and phosphatase inhibitor cocktail, for 1 hour at 4 °C and subsequently centrifuged at 15000 *g* for 10 min at 4 °C. Protein concentration was determined by Bradford method. The mitochondria soluble fraction (500 µg) was applied to a FPLC-Superdex S200 (30 × 1.5 cm) equilibrated with 20 mM Tris-HCl (pH 7.5), 10% glycerol, 5 mM NaCl, 10 mM β-mercaptoethanol, protease inhibitor cocktail and supplemented with 0.5 M NaCl. The eluted fractions were analyzed by Western blot using anti- ATP50/OSCP, anti-human TRAP1, and mouse monoclonal anti-Cyclophilin (ab110324) antibodies.

### Overlay assay

Purified F-ATP synthase, CyPD or TRAP1 were used. F-ATP synthase was purified from bovine heart as previously described [6]. Lauryl-maltose-neopentyl glycol (LMNG) stabilized F-ATP synthase complexes were eluted by a linear concentration gradient of 0–240 mM KCl in 40 mM HEPES pH 7.8, 150 mM sucrose, 2 mM MgCl2, 0.1 mM EDTA, 0.1 mM DTT, an d 0.05% (wt/vol) LMNG. F-ATP synthase fractions containing high amounts of native phospholipids were flash-frozen in aliquots of about 500 μl for later use, both in overlay assays and in electrophysiology experiments.The human CypD expression system was constructed by building the pSUMO- Cyclophilin D plasmid. The nucleotide sequence encoding for human CyPD was obtained from a Human cDNA placenta library (Clontech) by PCR. The resulting vector was used to transform *E. coli* strain BL21(DE3)pLysS and the correctness of the cDNA was confirmed by sequencing. Human TRAP1 was produced and purified as previously described [21]. Proteins were separated by SDS-PAGE and transferred onto nitrocellulose paper. The membrane was washed briefly with distillated water and incubated in overlay blocking buffer containing 50mM Tris-HCl pH 7.5, 200mM NaCl, 3% BSA and 0,1% polyethylene glycol for 1 h at room temperature. The membrane was then incubated with overlay buffer with or without 50 µg of purified TRAP1 or CyPD for 30 min. After 4 washing for 5 min with overlay buffer the membrane was incubated with or without TRAP1/CyPD antibody in the same buffer and bound TRAP1 or CyPD were detected by Western blot analyses.

### Blue native gel electrophoresis (BN-PAGE)

Experiments were carried out on isolated mitochondria to retrieve electron transport chain (ETC) complexes. Briefly, 500 µg of mitochondria were solubilized in 50 µl of extraction buffer containing 50 mM NaCl, 2 mM 6-aminocaproic acid, 1 mM EDTA and 50 mM imidazole/HCl pH 7.0, with different concentration of digitonin and immediately centrifuged at 100000 *g* for 25 min at 4 °C. Supernatants were supplemented with Coomassie Blue G-250 (Serva) and applied to 1D 3- 12% polyacrylamide gradient BNE (Invitrogen). After migration, the protein complexes were transferred on membrane and probed respectively with mouse monoclonal anti-ATPB (ab14730), anti SDHA (sc-166947) or anti-TRAP1 antibodies. In selected experiments, gels were stained with Coomassie Blue and monomer, dimer or oligomers of ATP-Synthase were cut and run on SDS-PAGE for protein identification by Western immunoblot or stained in-gel ATPase activity staining as reported [63] .

### Measurement of kinetic constant with surface plasmon resonance (SPR)

Recombinant mature human OSCP and CyPD were produced and purified in *E. coli* as described [21]. SPR measurements were performed with Biacore T100 biosensor system (GE Heathcare Bioscience) using a CM5 sensor chip with immobilized CyPD. CypD was immobilized on a CM5 utilising standard amine-coupling chemistry. The amount of CypD immobilized on the activated surface was typically between 800 and 1200 response units (RU). OSCP was injected at increasing concentrations. Unspecific binding and buffer interactions were subtracted from each sensorgram and the resulting curve were fitted using a Langmiur interaction model (ProteON Manager software, Bio-rad Laboratories, Hercules, CA) to obtain binding constant.

### Split GFP assay

sMPNST TRAP1 knock-out cells were grown on microscope slides and co-transfected with a combination of plasmids pcDNA3 mito-GFP1-9, pcDNA3-TRAP1-GFP10 and pcDNA3CyPD- GFP11, or pcDNA3ATP50-GFP11. Plasmids pcDNA3 mito-GFP1-9, pcDNA3-TRAP1-GFP10 and pcDNA3ATP50-GFP10 were previously described [21]; pcDNA3CyPD-GFP11 and pcDNA3ATP50-GFP11 plasmids encoding for human CyPD and OSCP respectively, were modified with the eleventh b strand of sfGFP at the C terminus (EFSGSGGGSGGGSTSEKRDHMVLLEYVTAAGITDAS). Cells were stained with 50 nm MitoTracker red for 30 min, 48 h after transfection, to detect mitochondrial network, fixed with 4% PFA and visualized with a LSM 700 confocal microscope (Zeiss, Oberkochen, Germany). For evaluating the interaction between CyPD and OSCP, sMPNST cells were transfected with the pcDNA3 mito-GFP1-9, pcDNA3CyPD-GFP11 and pcDNA3ATP50-GFP10 plasmids, stained, fixed, visualized with a fluorescence Leica DMI600B microscope (20X objective) and analyzed using LAS AF (Leica Microsystems, Wetzlar, Germany) and ImageJ^®^ software (National Institutes of Health, University of Wisconsin, WI). The percentage of cells showing the interaction by reconstitution of GFP was calculated as the ratio between green (GFP cells) and red (total number) cells; at least 2500 cells were analyzed for each condition.

### Nonyl Acridine Orange (NAO) staining

Cells, were seeded at a density of 5 x 10^4^ cells/well in a 12-wells tissue culture plate. After 24 h cells were stained for 30 min at 37 °C with NAO dye (200 nM, Merck), washed, detached with trypsin, centrifuged at 1000 *g* for 5 min, and suspended in the saline buffer. NAO staining was assessed by flow cytometry using the FACS Canto II flow cytometer (Becton Dickingson).

### ATP hydrolysis

ATPase activity was measured monitoring the rate of NADH oxidation in 4 x 10^6^ permeabilized cells or in 50 μg of pig heart MgATP-submitochondrial particles (SMP) in ATP regenerating buffer [11]. The buffer for cell samples contained 50 mM KCl, 50 mM Tris-HCl, 30 mM sucrose, 4 mM MgCl_2_, 2 mM EGTA adjusted to pH 7.4, 4 units/ml pyruvate kinase, 3 units/ml lactate dehydrogenase, 4 mM phosphoenolpyruvate, 2 mM ATP, 0.2 mM NADH. When cells were analyzed, 10 μM alamethicin and 10 μM sodium decavanadate were added. NADH absorbance was measured spectrophotometrically at 340 nm, 37°C. Inhibition of Mg^2+^-ATPase activity was obtained by adding 1 μM oligomycin. Values were normalized for mg of proteins.

### Calcein staining

Cells were seeded and cultured for 2 days in coverslips (10^5^/sample) with normal culture media. The coverslips were then transferred into an open chamber and incubated in HBSS without phenol red supplemented with 8 mM CoCl_2_ and 0,8 μM CsH for 10 minutes and for further 10 minutes with 0.5 μM Calcein-AM with or without 2 μM CsA. After calcein-AM incubation, cells were washed with PBS, incubated with HBSS in the presence of CsH and analyzed with a fluorescence microscope (Leica DMI600B). After 1 minute, the Ca^2+^ ionophore Calcimycin A23187 (Sigma) was added; 1 picture was recorded every 30 sec for 25 min.

### Evaluation of mitochondrial membrane potential by microscopy

Mitochondrial membrane potential was tested using the fluorescent potentiometric compound tetramethylrhodamine methyl ester (TMRM, 20 nM; Invitrogen). Cells were incubated in DMEM without phenol red with 0.1% FBS and CsH (1 µM) to inhibit P-glycoproteins. Recordings were performed with a fluorescence microscope (Leica DMI600B) and analyzed using LAS AF software (Leica) as described [64].

### Electrophysiology

Electrophysiological properties of F-ATP synthase were assessed by single-channel recording following protein insertion into artificial planar lipid bilayers by direct addition of 8 μg of purified protein. Membranes were prepared by painting a solution of soybean asolectin (10 mg/ml in decane, Sigma) across a 250 μm-diameter hole on a Teflon partition separating two compartments filled with a recording solution (150 mM KCl, 10 mM HEPES, pH 7.4) before membrane painting. The two compartments are defined as *cis* and *trans*, and all voltages refer to the *cis* side, zero being the *trans* (grounded) one. Currents were considered as positive when carried by cations flowing from the *cis* to the *trans* compartment. Membrane capacity ranged from 50 to 150 pF (average 100 pF) and no current leakage was detectable. F-ATP synthase was directly added to the recording chamber followed by the additions specified in the text; TRAP1 and CyPD were directly added in the recording chamber in the molar ratios to F-ATP synthase specified in the text. Data were acquired at 10 kHz through a Bilayer Clamp BC-525C amplifier (Warner Instruments, Harvard Bioscience, Inc.) and low-pass filtered at 500 Hz. Data were digitized with a Digidata 1322 A interface and pClamp software suite (all from Molecular Devices) and analyzed offline using the same software. Conductance values (G) were calculated from stable current signals corresponding to open states on the basis of Ohm’s law. Maximal conductance (Gmax) was calculated from the maximal stable current level (i.e., events lasting at least 10 ms) in the recording interval.

### Statistical analysis

Data were analyzed and presented as mean ± standard error of the mean (SEM) in all figures. Pairs of data groups were analyzed using paired and unpaired two-tailed Student’s t tests. Statistical significance was determined using Origin^®^ 8 (OriginLab, Northampton, MA). Protein quantification and assessment of significance for the mass identification was determined using Prism 6.0 with Benjamini-Hochberg procedure correction. Results with a *p* value lower than 0.05 were considered significant; \*\*\**p* < 0.001, \*\**p* < 0.01, \**p* < 0.05 compared to controls. Each experiment was repeated at least three times.

## Acknowledgements

We thank Elena Trevisan for invaluable technical assistance. A.R. and P.B. were supported by Associazione Italiana Ricerca Cancro (AIRC grant IG 2017/20749 and IG 2019/23129, respectively). F.C. was recipient of a Young Investigator Award Grant from Children’s Tumor Foundation.

## Author contributions

G.C.: conceptualization, visualization, methodology, investigation, formal analysis, writing-original draft; A.U.: investigation, formal analysis; M.G.: methodology, formal analysis; M.V.: investigation, formal analysis; A.N.: methodology, investigation, formal analysis; A.F.: investigation, formal analysis; F.C.: investigation, formal analysis; I.M.: conceptualization, formal analysis; C.G.: methodology, resources; E.T.: investigation, formal analysis; A.M.B.: methodology, formal analysis; G.L.: conceptualization, methodology, visualization, formal analysis; P.B.: conceptualization, review and editing original draft, funding acquisition; A.R.: conceptualization, writing original draft and subsequent review and editing of the text, funding acquisition, project administration, supervision.

